# Detecting significantly recurrent genomic connections from simple and complex rearrangements in the cancer genome

**DOI:** 10.1101/2023.10.13.561748

**Authors:** Shu Zhang, Kiran H. Kumar, Ofer Shapira, Xiaotong Yao, Jeremiah Wala, Frank Dubois, Rose Gold, James E. Haber, Andrew Cherniack, Marcin Imielinski, Simona Dalin, Rameen Beroukhim

## Abstract

The detection of somatic genetic alterations that recur across cancer genomes more than expected by chance has been a major goal of cancer genomics, as these alterations are enriched for “driver” events that promote cancer. Multiple methods have been developed to detect driver point mutations and copy-number variants, but methods to detect driver rearrangements have largely not been pursued. Unlike point mutations and copy-number alterations, which can be assigned to a single genomic locus, rearrangements connect two distant genomic loci, and possibly more in the case of complex or clustered events. Here, we explore genomic features that predict the rate at which any pair of loci will be connected by rearrangements and describe two methods to detect rearrangements that recur more often than this background rate. The first, SVSig-2D, detects pairs of loci that are directly connected by a single rearrangement; the second, SVSig-2Dc also detects loci that are recurrently connected indirectly through two or more rearrangements. When applied to a pan-cancer dataset of over 2,500 cancers, these methods identified 80 significantly recurrent simple rearrangements and 29 complex rearrangements, including both known and novel events. Intriguingly, though both recurrent simple and complex rearrangements tended to be tissue-specific, this was less true for the complex events. The detection of recurrent rearrangements with methods such as these will be an essential component of cancer genomics in the whole-genome sequencing era.

## Introduction

Somatic rearrangements can have major impacts on cancer genomes. These events connect distant genomic loci, bringing originally distant genetic elements (e.g. genes and regulatory regions) into direct apposition and often amplifying or deleting vast tracts of the genome. As such, understanding their genomic impact is essential to cancer genome analysis.

A major goal of cancer genome analysis is to distinguish between “driver” events that promote cancer development and progression and “passenger” events that do not. In the case of rearrangements, driver events can promote cancer through the copy-number alterations (SCNAs) they create: amplifying oncogenes or deleting tumor suppressors. However, they can also drive cancer through the novel connections they create, leading to new fusion genes and hijacking of regulatory elements (e.g. enhancer hijacking [1,2] that profoundly alters gene expression). Examples of these types of driver rearrangements include fusions of BCR and ABL in chronic myelogenous leukemias, TMPRSS2 and ERG in prostate cancer, and IGH and MYC in Burkitt’s lymphoma [3–5].

Methods to detect driver genetic alterations in cancer often rely on assessing their recurrence above randomly generated “background” alterations. Due to selection biases (all cancers have driver events), driver events tend to recur across cancers more frequently than passengers. Based upon this principle, multiple methods have been developed to detect recurrence of SCNAs, single nucleotide variants, and small insertion-deletion events (indels) [6][7][8]. These have led to the discovery of new oncogenes and tumor suppressor genes, thereby improving our understanding of cancer evolution and indicating promising therapeutic strategies.

However, there are no widely used methods to detect recurrent novel connections generated by rearrangements. The reasons for this are three-fold. First, the vast majority of rearrangements connect intronic and intergenic loci, thus making it unclear which genes and regulatory elements they are disrupting. Second, rearrangements are fundamentally different from other genetic events in that they connect two or more genomic loci, as opposed to altering a single locus. Thus, detecting recurrent connections requires a two-dimensional approach rather than detecting recurrent events along a one-dimensional genome. Third, these connections can reflect multiple neighboring rearrangements in a single complex event, such that the functional effects of one rearrangement are modulated by its neighbors. Indeed, more rearrangements exist within complex events than as isolated simple events, and are evident in phenomena such as chromothripsis or chromoplexy[9,10].

Detecting driver events that recur above background rates of genetic alteration requires accurate models of those background rates. Rearrangement densities can vary with their locations within the genome, distance between breakpoints, and underlying DNA damage repair mechanisms from which they were formed, as well as many other variables [11–13]. Thus, to develop methods that accurately detect driver rearrangements, the effects of these factors on rearrangement densities need to be understood.

Here, we present SVSig-2D and SVSig-2Dc, two methods that utilize the two-dimensional breakpoint distributions of rearrangements to identify positively selected events in cancer. In both models, we employed the empirical patterns of rearrangement features to calculate background rates of rearrangement formation. SVSig-2D considers each SV independently of all others, while SVSig-2Dc accounts for novel connections generated by two or more neighboring rearrangements. We measure the effects of a variety of underlying variables on rearrangement rates through analysis of nearly 300,000 SVs from the PCAWG Consortium and apply SVSig-2D and SVSig-2Dc to this dataset to detect known and novel recurrent rearrangements.

## Results

### Overview of Method

We took a systematic approach to detect recurrent rearrangements in cancer genomes, searching for pairs of genomic loci that connect to each other more often than expected due to mechanistic factors. Since we were interested in finding recurrently connecting pairs of loci, we used a two-dimensional strategy. Specifically, we created an adjacency matrix with each dimension representing the genome, divided into 500 kb bins, and each element (or “tile”) representing the number of rearrangements connecting the two bins (**Figure 1a**). We chose a bin size of 500 kb to approximate the spans of topologically associated domains (TADs) [14].

**Figure 1.**
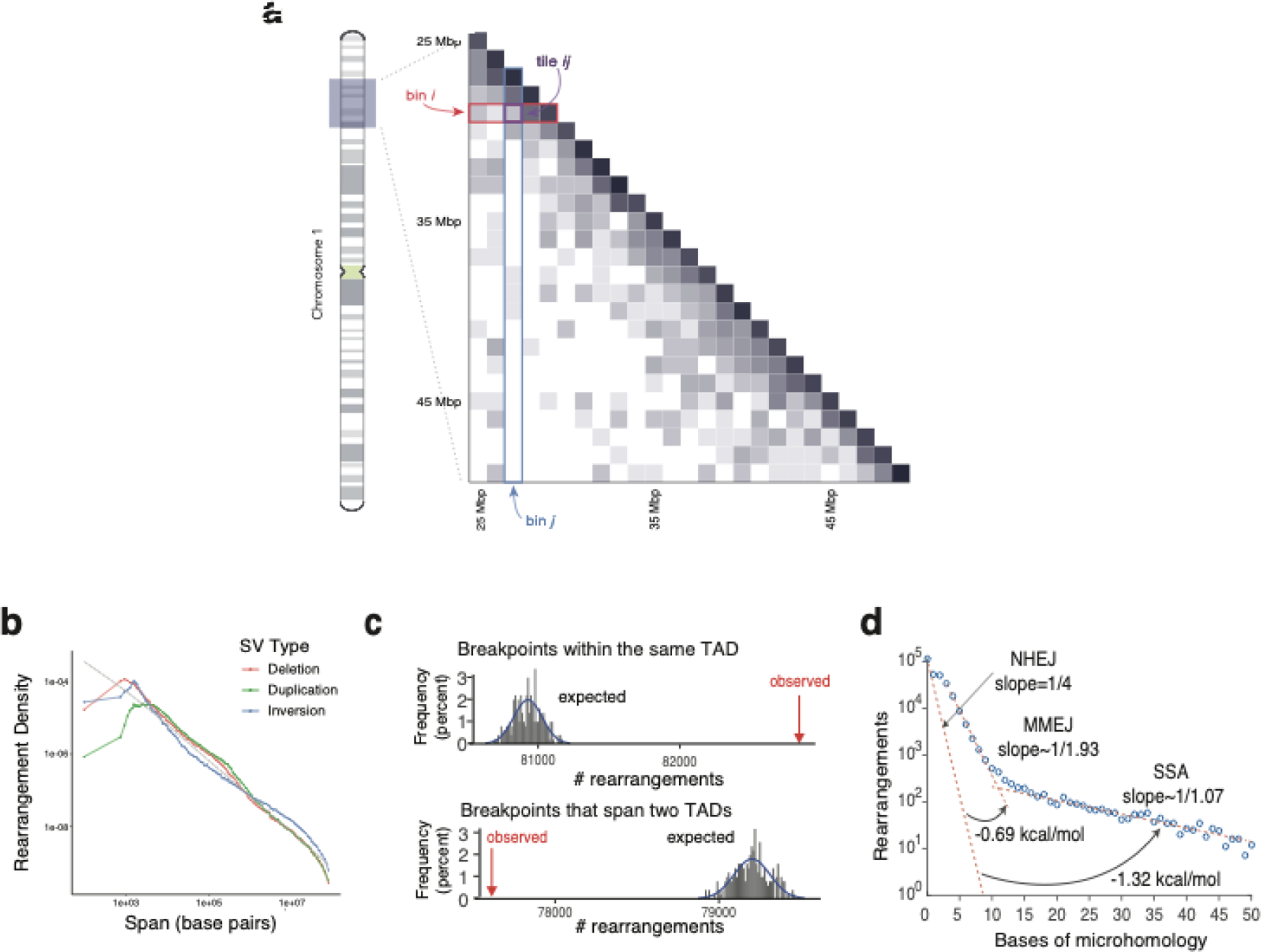
Predictors of rearrangement formation genome-wide. a) Representation of the genome as a 2D adjacency matrix. The linear genome is binned into 1 Mbp bins. The intersection of two bins, on axes i and j, is represented by a tile *ij*. b) Distribution of intrachromosomal rearrangement spans, grouped by rearrangement type. Spans approximately follow a 1/span distribution, represented by the dashed line. Rearrangements less than 200 bp and greater than 100 Mbp were excluded. c) Observed and expected number of rearrangement breakpoints that fall either within a TAD or across two TADs. The distribution of the expected number of breakpoints is determined by permuting breakpoints across the genome. d) Distribution of microhomology length at rearrangement breakpoints. The curve appears to divide into three components with different slopes, with microhomology lengths corresponding to known mechanisms of NHEJ, MMEJ, and SSA. The deviation of these slopes from the expected ¼ decline for random connections indicates the energetic favorability of each additional base of homology (adapted from Figure 5d, Li et al, 2020).

We wanted to identify rearrangements that underwent positive selection in cancer genomes under the hypothesis that they were causal. Therefore, we considered genome-wide patterns that shape the distribution of rearrangements, and created a background model representing how often rearrangements should form by chance alone. We then compared this model to the observed data to detect adjacencies that recur significantly more often than predicted, thereby representing candidates for undergoing positive selection (i.e. “driver” rearrangements).

### Biological Factors That Shape Connected Loci

In order to develop the background model we identified likely mechanistic biases that shape the distribution of rearrangements--and specifically the pairs of loci that get juxtaposed--across the genome. We analyzed 292,284 rearrangements detected among 2,693 cancers spanning 30 histological types from the International Cancer Genome Consortium (ICGC) Pan-Cancer Analysis of Whole Genomes (PCAWG) effort [8,15,16]. We considered four biological determinants of rates at which loci are juxtaposed: the genomic distance between these loci in the reference genome (termed the rearrangement “span”); the extent of homology between them; the one-dimensional rate at which each locus is involved in rearrangements; and other characteristics of these loci, such as the types of sequences they involve, their replication timing, and epigenetic characteristics.

The dominant feature determining the rates of rearrangement between each pair of genomic loci appeared to be the distance between them. Hi-C data have shown that, broadly speaking, DNA is packed in a fractal globule structure in which 3D distances between genomic loci scale as inversely proportional to the 1D genomic distance between them. Likewise, rearrangement frequency followed a 1/span distribution, as previously noted [15,17,18]). The 1/span distribution was largely maintained across tissue types and rearrangement topologies (**Figure 1b, Supplementary Figure 1**a). Deletions and duplications differed from the 1/span distribution respectively by only 19% and 26% on average across all spans, with a slight excess of short events; however inversions differed by 42%, with an excess of longer rearrangements. The cancer types that deviated most from a 1/span distribution were kidney carcinomas (KIRC) and sarcomas (SARC), which favored longer events, and oral cancers (ORCA) and endometrial cancers (UCEC), which favored shorter ones. At higher resolution, the fractal globule 3D structure of DNA is modulated by the presence of topologically associated domains (TADs); 3D interactions are more common between DNA loci within a TAD than between TADs. Accordingly, we found that rearrangements were more likely to occur within TADs than between TADs. (**Figure 1c**). These results are in line with prior studies of SCNAs [17–20] and rearrangements in smaller cohorts [21,22].

We also found that variations in genomic sequence and epigenetic features, including presence of LTR or LINE sequences and heterochromatin, influence which locus pairs become juxtaposed. Across 45 combinations of 10 sequence features (**Supplementary Figure 1**b), 14 were significantly associated with connection frequency [15]. Among these, LTR-LTR connections were most significantly enriched.

### Rearrangement Sequence Homology and DNA Damage Repair

Connected loci also tend to have higher levels of homology than expected by chance, reflecting processes of DNA damage repair that helped generate them. Pairs of loci with little or no homology are joined by non-homologous end-joining (NHEJ), whereas microhomology-mediated end-joining (MMEJ), single-strand annealing (SSA), and homologous recombination (HR) all connect loci with higher levels of sequence homology [23]. If rearrangements were generated only by NHEJ, we would expect a four-fold drop in the number of rearrangements with each increasing base of microhomology, corresponding to the probability of encountering one of the four DNA nucleotides. In contrast, we observed an approximately two-fold drop per base for rearrangements with junction microhomology between 3 and 10 bp, and only a 7% decrease in the number of rearrangements observed with each increasing base of homology past 11 bp (**Figure 1d**).

These results are consistent with NHEJ being responsible for 70% of somatic rearrangements (as previously reported ([15]), and also suggest a substantial and quantifiable energy benefit with increasing homology for MMEJ and SSA. MMEJ has previously been found to apply to sequences with more than two bases of homology; the slope of the drop in this range (**Figure 1d**) suggests that each additional base of homology provides a 0.69 kcal/mole benefit for MMEJ (see Methods). SSA has been found to require 13 or more bases of microhomology in yeast ([24]), and the slope of the drop in this range suggests that each additional base of homology provides SSA with a 1.32 kcal/mol energy benefit. In addition, the transition to SSA appears to occur at lower levels of microhomology in human cancer (11 bp) than in yeast.

We further investigated the relationship between mutations in cancer driver genes and distribution of usage of each type of DSB repair. We assigned rearrangements with 0 bases of microhomology (39% of rearrangements) to NHEJ, 4 - 9 bases (12%) to MMEJ, and 13 or more bases (1%) to SSA. We considered rearrangements with 1-3 and 10-12 bases of homology to be ambiguous and therefore excluded them.

We first looked for patterns of estimated DNA repair usage and rearrangement frequencies across our cohort by clustering samples according to the number of rearrangements in each tumor and the fractions of these that were attributed to each type of DSB Repair.

The tumors segregated into four clusters: two clusters with, respectively, small and large numbers of rearrangements attributed to NHEJ, a cluster with a high rearrangement burden overall, and a cluster with near-average values for both the number of rearrangements per sample and the fractions assigned to NHEJ, MMEJ, and SSA (**Figure 2a**, **Supplementary Table 1**). Only 45% of rearrangements in the “NHEJ low” cluster were attributed to NHEJ whereas 54% were attributed to MMEJ. Conversely, 84% of rearrangements in the “NHEJ high” cluster were attributed to NHEJ. Tumors in the “High Rearrangement Burden” had an average of 635 rearrangements, whereas the “Average” cluster had a mean of 124 rearrangements. The vast majority of tumors segregated into the “Average” (48%) and “NHEJ High” (41%) clusters; only 2% and 8% of tumors segregated into “NHEJ low” and “High Rearrangement Burden” respectively.

**Figure 2.**
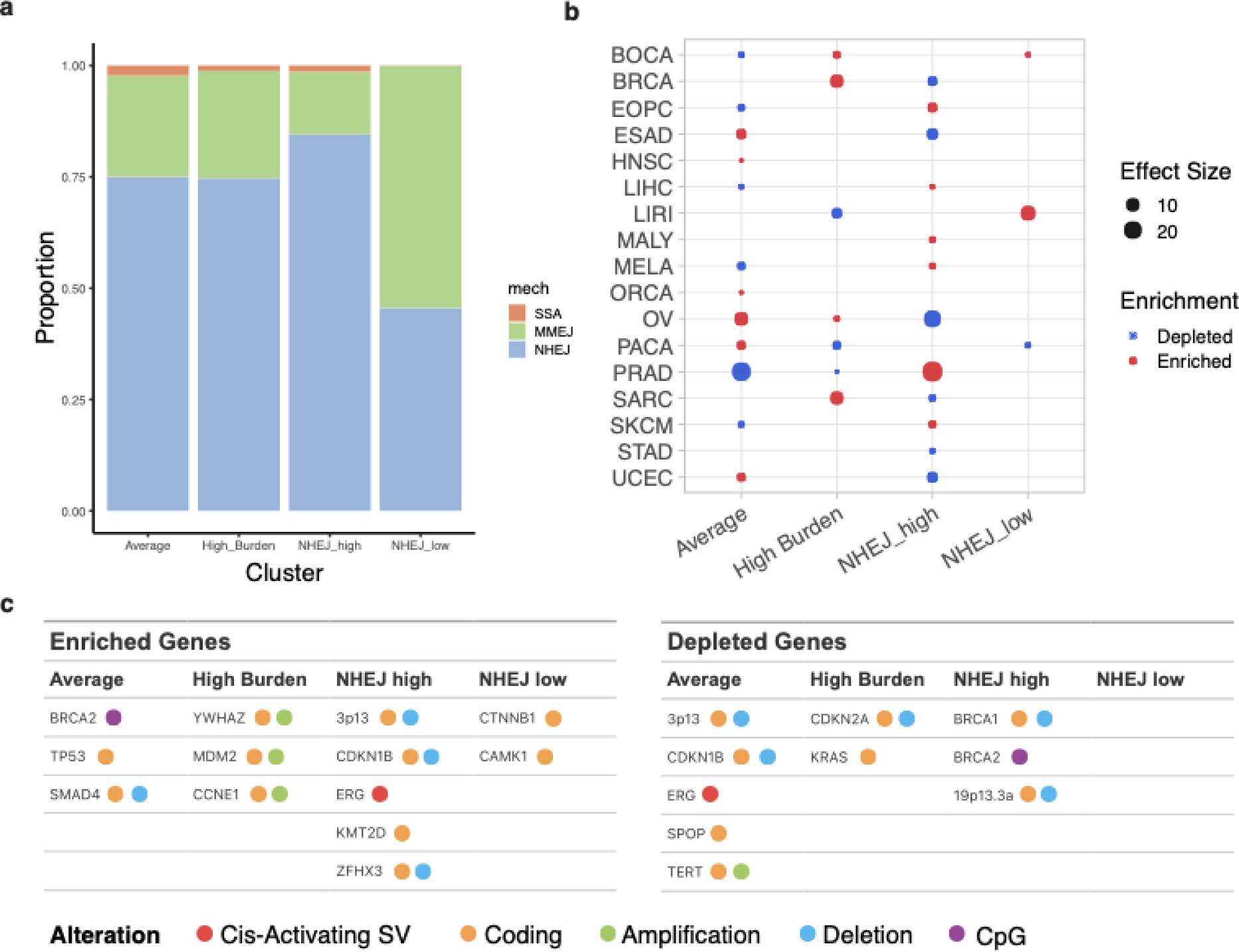
Rearrangements cluster according to the DNA damage repair mechanisms that formed them. a) K-means clustering of samples based on the proportion of DNA damage repair mechanisms that formed their rearrangements. DDR mechanisms were assigned based on the number of bases of microhomology present at the rearrangements’ breakpoints. b) Tissue enrichment across clusters. For each cancer type on the vertical axes, a Fisher’s exact test was performed, followed by FDR correction. The size of the circle represents the Fisher’s exact test statistic, which we use as the effect size . c) Enrichment of known driver genes across clusters. Fisher’s exact tests were performed for each known driver. Driver genes are colored by the means by which the gene is altered.

Both cancer types and driver alterations were non-uniformly distributed across these clusters (**Figure 2b-c**) (**Supplementary Tables 1-2**). The “Rearrangement Burden High’’ cluster was enriched for cancers designated as breast, bone, ovarian, and sarcoma (which itself included sarcomas of bone). All of these cancer types are known to have large SCNA burdens due to frequent TP53 inactivation in breast and ovarian cancers and frequent ring chromosome formation in sarcomas leading to high-level amplifications of several genes including MDM2 [25,26]. The “NHEJ high” cluster was enriched for melanomas and depleted of gynecological cancers and cancers of the gastrointestinal tract. Melanomas are radiation-resistant, which is consistent with increased NHEJ [27,28]. The “average” cluster had the opposite representation of cancer types. Finally, the “NHEJ low” cluster was enriched in liver and bone cancers and depleted in pancreatic adenocarcinomas.

The clustering was robust. Within the dataset, six cancer types were independently profiled by two or three projects conducted in different countries. In four cases, tumors profiled by at least one of these projects were significantly (p<0.05) enriched or depleted in at least one cluster. In all four of these cases, at least one of the other projects was also enriched in at least one of those same clusters (p<0.0001)--indicating the reproducibility of these results. In the “Average” cluster, prostate cancers from three different countries (US, UK, AU) and ovarian cancers from the US and Australia clustered together. In the “Rearrangement Burden High” cluster, breast cancers from the US and UK and pancreatic cancers from Canada and Australia clustered together. Finally, in the “NHEJ high” cluster, prostate cancers from US and Canada, ovarian cancers from Australia and the US, and breast cancers from the US and UK clustered together (**Supplementary Table 3**).

Finally, we tested to see which cancer driver gene mutations were enriched in the four clusters. Similar to the results of the tissue type enrichment analysis, the “Average” cluster and the “NHEJ high” cluster were anticorrelated in terms of enriched and depleted genes. The “Average” cluster was enriched in mutations in BRCA2 CpG sites, potentially altering methylation patterns at BRCA2 CpG islands. The “NHEJ low” cluster was enriched in CTNNB1 and CAMK1 coding mutations, and the “Rearrangement Burden High” cluster was enriched in YWHAZ, MDM2, and CCNE1 coding amplifications and depleted in CDKN2A coding deletions and KRAS coding mutations. These genes affect G1/S checkpoint regulation; we hypothesize that tumors with these mutations enter S phase too early, leading to instability and accounting for their high rearrangement burden.

We also found 116 genes whose mutation status was significantly associated with the distribution of rearrangement homology levels. These include genes known to be involved in DNA damage repair (e.g. MGMT, BRCA1, FANCD2) and genes that are not known to be involved (e.g. APOB, KRAS, CDKN1A). These findings suggest undiscovered connections to DNA damage repair.

### The predictive value of rearrangement features on connection frequency

We next evaluated how strongly these determinants of connection frequency predicted rearrangement rates, in two ways: First, we calculated the difference between models that alternatively accounted for or did not account for each determinant. Second, we calculated the predictive value of each determinant using ρ^2^ values from Spearman correlations between the determinant and connection rates.

The rearrangement span was by far the most important, followed by levels of microhomology and the total predictive value of all combinations of sequence features indicated above (**Supplementary Figure 1**b). The effect of accounting for genomic distances between loci was 2630 times greater than the effect of accounting for breakpoint homologies and 1.2 x 10^11^ times greater than the effect of accounting for the most significant association with combinations of sequence features: LTR-LTR connections.

The result of the dominating effect of span is that at long spans, few rearrangements are observed and the pattern of rearrangements is largely stochastic. We determined the extent to which rearrangement rates determined from one-half of the dataset predicted rearrangement rates in the other half. Specifically, we divided the genome into 500 kb bins, and considered the rates at which rearrangements connected any two bins (a “tile”) (**Figure 1a**, see Methods). At short spans (<1 Mb), the predictive value, or correlation, between rearrangement rates in each tile was high between the two halves of the data: ρ=0.7. At longer spans and for interchromosomal events, however, stochasticity dominated: most tiles contained no rearrangements, and only a few contained one or two. As a result, the correlation between the two halves of the dataset was actually negative (ρ=-0.28 and -0.8, respectively)--reflecting the sparsity of the data. At rearrangement spans of less than 1 Mb, where the data were dense enough to determine associations, span appeared to predict over 30% of the variation in rearrangement rates (ρ=0.56). The next best predictor was 1D breakpoint density, which predicted 3% of the variation in rearrangement rates (ρ=0.18).

We wanted to see if our initial predictors of rearrangement formation held similar relationships to rearrangement frequency once rearrangements were discretized into the adjacency matrix (**Figure 1a**). Using 500kb bins, we found that breakpoint density had the largest correlation with connection frequency across short tiles, with ρ=0.53 (Spearman, p<1e-150). These correlations were lower across long and interchromosomal tiles, with ρ = 0.12 and 0.03, respectively (Spearman, p<1e-150 in both cases). Rearrangement span was the largest predictor across long (>1Mb) intrachromosomal tiles, with ρ = 0.27. Short tiles were excluded from this analysis since there was minimal variation in the average span across these tiles. Finally, LTR, LINE, and SINE regions were found to have little association with the frequency of rearrangements that spanned those regions, with ρ < 0.01 for all tiles and sequence features except for short tiles and SINE regions, which had a modest correlation of ρ = 0.2 (Spearman, 1.01e-52). We did not compare microhomologies per tile with SV rates because each tile represents 250 billion possible SVs, each with its own microhomology.

### Developing a Background Model for Simple Connections

We next developed an approach to detecting significantly recurrent connections, which we call SVSig-2D, comprising two major steps. *First*, SVSig-2D estimates the background probability *p_ij_*with which we expect any two loci *i* and *j* to combine, in the absence of selective pressure. We based this background model upon the one-dimensional breakage rates across the genome and the distance between *i* and *j*, because of the dominant effects of these relative to other predictors of rearrangement frequencies (**Figure 3a**). SVSig-2D calculates the probability of rearrangements occurring within a 2D tile as the sum of *p_ij_* over the *ij* pairs represented in the tile. *Second*, SVSig-2D determines which tiles have more rearrangements than would be expected according to this background model. Here, SVSig-2D calculates a one-sided p-value, using a binomial (N, *p_ij_*) where N is the total number of rearrangements in the dataset. These p-values are then converted to FDR-corrected q-values to account for multiple hypotheses. Here we consider tiles with q-values of less than 0.1 to contain candidate driver rearrangements.

**Figure 3.**
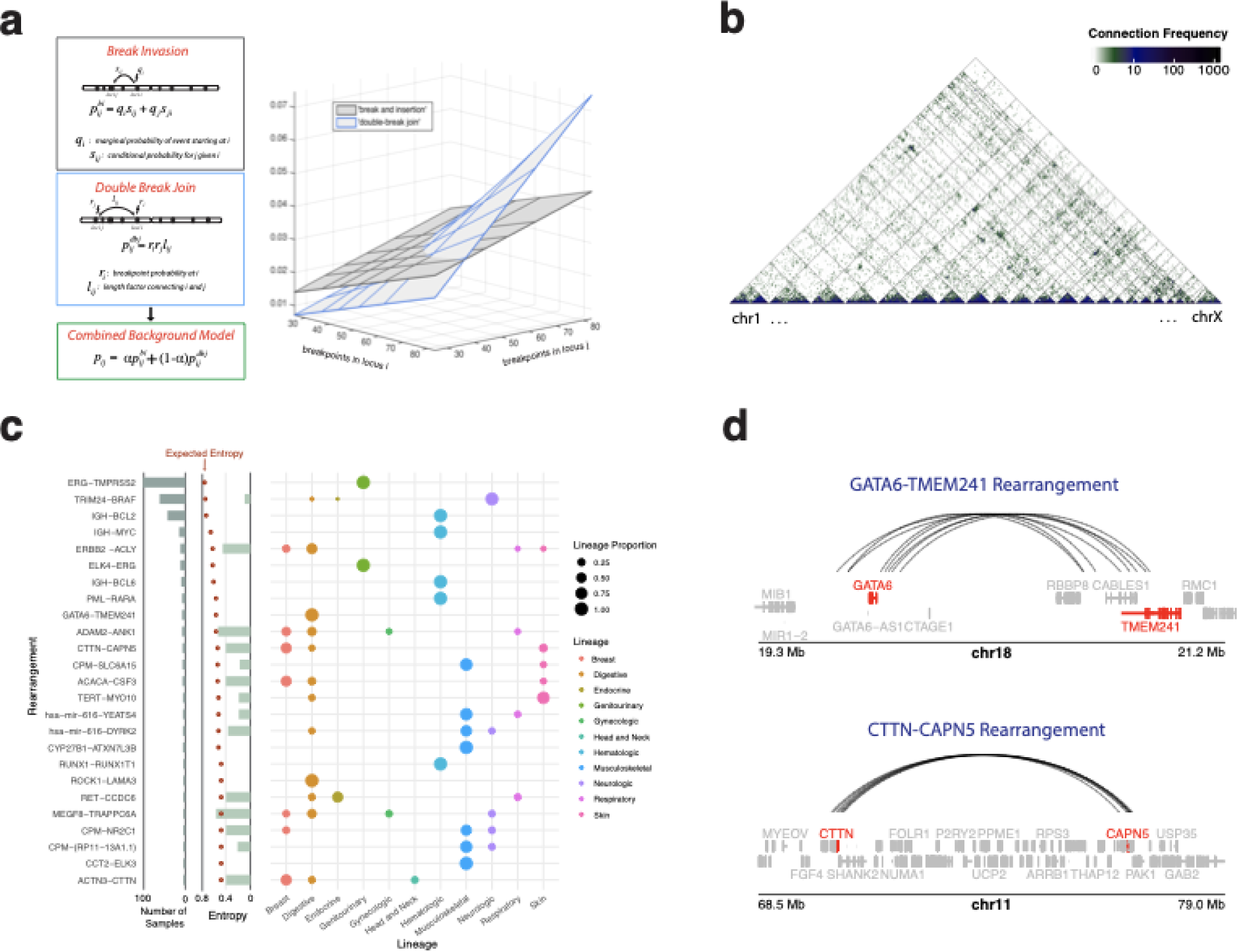
SVSig-2D identifies significantly recurrent rearrangements. a) The break-invasion and double-break join background models, and the linear combination used by SVSig-2D. The variation of each model with breakpoint rates is shown on the right. b) Observed connection frequencies genome-wide, represented on a log10 scale. c) The 25 most statistically significant events identified by SVSig-2D. For each event, the proportion of affected samples that fall under each lineage is shown by the size of the circle. Entropy values represent levels of lineage specificity. The number of samples containing each rearrangement is also indicated. d) Novel significant rearrangements involving GATA6-TMEM241 and CTTN-CAPN5. Eight samples from digestive lineages contained the GATA6-TMEM241 rearrangement. Seven samples, four of which were breast cancers, contained the CTTN-CAPN5 rearrangement.

To calculate the background rates *p_ij_*, we considered the two simplest possible models. Because we cannot distinguish between i joined to j and j joined to i, we recognized that p_ij_ should be a symmetric matrix. The first is based on the additive probabilities of breakage at each locus:

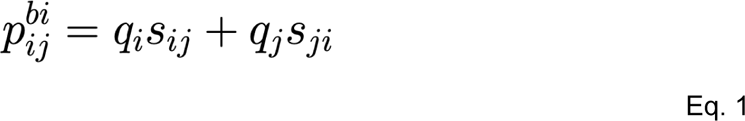

where *q_i_* is the marginal probability of a rearrangement initiated in locus *i* and *s_ij_* is the conditional probability that a break at *i* will connect to site *j*. This “break-invasion” model is reminiscent of mechanisms like non-allelic homologous recombination (NAHR), which involve a break in one locus followed by invasion into another[29]. The second model is based on multiplicative probabilities:

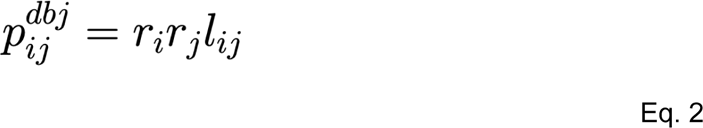

where *r_i_, r_j_* are the breakpoint densities and *l_ij_* is a length factor connecting *i* and *j*. This “double-break join” model is reminiscent of non-homologous or microhomology-mediated end joining (NHEJ or MMEJ [30]), which involve separate breaks in two loci that are erroneously joined.

The extent to which different classes of rearrangements fit either model is partially indicative of the physical process that generated those rearrangements. We tested rearrangements stratified by level of homology, topology, and span for their fit to each model. Short rearrangements tend to reflect an even mixture of the two models, whereas long and interchromosomal rearrangements favor break-invasion. Increased homology is associated with only a slight preference for break-invasion, primarily among interchromosomal events. Rearrangements whose junctions included an insertion longer than 10 bp (independent of junction homology), a characteristic often attributed to microhomology-mediated break-induced replication (MMBIR) [31], were also 10% more likely to fit the break-invasion model (p<10^-4^). Simple rearrangements tended to fit the break-invasion model whereas complex events tended to fit the double-break join model (p<10^-4^). And while rearrangements of all lengths tended to fit the break-invasion model, interchromosomal rearrangements were more likely to fit the double-break join model than intrachromosomal rearrangements.

In every case, the observed rearrangements appeared to reflect a mixture of both models. The SVSig2D background model is therefore a weighted sum of these two underlying models:

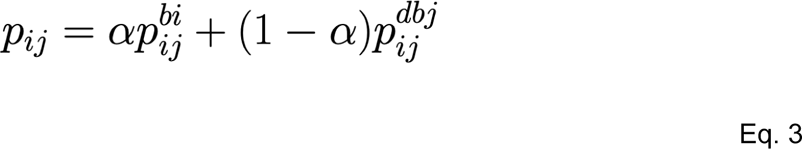

with α set to fit the observed data most closely. This α varied among rearrangements of different spans, so we use different values for short (500 kb to 1 Mb), long (greater than 1 Mb), and interchromosomal events. Our estimates of the span distribution were less accurate at spans that were less than the size of a bin (500 kb), so we exclude these extremely short events here. SVSig-2D then calculated the p-value representing enrichment of events in each tile using the binomial (N, p_ij_), where N represents the total number of events in the dataset.

In addition to detecting tiles that were enriched for rearrangements, we wanted to consider the extent to which those rearrangements clustered within the tile. SVSig-2D therefore calculates a second p-value using a t-test comparing the average 2D distance between all events in a tile to the distribution of average 2D event distances for random data. These two p-values are then combined using Fisher’s method to detect recurrent, clustered connections, and corrected for multiple hypotheses using the False Discovery Rate (FDR) method [32].

#### Testing the Background Model

We next tested this approach with simulated data, specifically asking two questions: *first*, how accurately does SVSig-2D determine α under different ground truths, and *second*, what was its propensity for false positive results?

To test how accurately SVSig-2D determines α, we randomly generated sets of SVs following various mixtures of the two background models of rearrangement formation (break-invasion and double-break join). When the number of SVs we used approximated that of the PCAWG dataset, we found that SVSig-2D accurately recaptured α to within 0.13 at all SV spans (**Supplementary Figure 2**a**, Supplementary Table 4**). As expected, our accuracy in determining α depended on the number of rearrangements generated per dataset **(Supplementary Table 4**). For datasets as small as 50,000 SVs, SVSig-2D recaptured α to within 0.3 units of its true value.

To test the propensity for SVSig-2D to generate false positive results, we applied it to simulated data. SVs were simulated without positive selection while maintaining the span distribution and marginal breakpoint densities from the PCAWG dataset. In this case, SVSig-2D only detected significantly recurrent connections in 1/10 independently simulated datasets. We further examined the extent to which incorrect estimates of background connection rates would lead SVSig-2D to generate false positive results. First, we tested the span distribution by raising the distribution to a range of powers, where powers greater or less than 1 steepen or flatten the distribution, respectively. The model produced false positive events when the span distribution was raised to a power less than 0.3 or greater than 1.13 (**Supplementary Figure 2**b). At the largest spans (∼100,000,000 bp), this corresponds to 444 times larger and 0.29 times smaller approximations of the span distribution, respectively. The model is more sensitive to incorrectly estimating low background rates at higher spans. We similarly tested the marginal breakpoint densities, and an exponential factor less than 0.64 and greater than 1.92 produced false positive results.

### Recurrent Connections detected by SVSig-2D

We found 80 recurrent connections, including 15 known oncogenic rearrangements (as described in the COSMIC database) and 65 that have not been shown to be oncogenic (Supplementary Table 5). Among the 65 novel events, 25 included breakpoints spanning at least one cancer gene. Furthermore, 25 of these connections (including 19 novel ones) were found in 5 or more unique samples. Seven connections (all known oncogenic rearrangements) were found in 10 or more samples. As in a prior study (Rheinbay et al, Nature 2020), we found that recurrent connections tend to be enriched in specific lineages. We grouped cancer subtypes by the overarching body systems they belonged to, following the NCI and ICD10 designations (**Supplementary Table 6**). Of all recurrent event clusters, 72.5% had a majority of samples derived from one lineage, including 66% of clusters that were not known to be oncogenic. This was significantly higher than if lineages were distributed randomly (**Supplementary Figure 3**a, permutation test, p<1e-4 in both cases), where only an average of 18.73% of events would have a prevailing lineage. We also measured the within-cluster randomness of the distributions of lineages with an entropy score. Of the top 25 events, every event except one had lower entropy than expected for a total of 63% less entropy than expected (**Figure 3c**). We conclude that recurrent connections tend to be lineage-specific.

Our analysis differs from Rheinbay et al in the methods used to bin the genome and estimate background rearrangement rates. This resulted in the detection of several new recurrent connections that involve genes that are not known to be cancer drivers but have been connected to cancer. For example, we observed a *GATA6*-*TMEM241* rearrangement in eight samples from colon adenocarcinoma, esophageal, gastric, and pancreatic cancers (**Figure 3d**). Only one of these appears to alter the *GATA6* coding sequence, and GATA6 rearrangements were associated with a 1.73-fold increase in GATA6 expression compared to samples without GATA6 rearrangements in the same lineage (**Supplementary Figure 3**b; p=0.06). GATA6 is a transcriptional regulator that plays a role in gut development and has been suggested to have tumor suppressive effects in pancreatic cancer and oncogenic effects in esophageal adenocarcinoma [33,34]. Similarly, a *CTTN*-*CAPN5* rearrangement occurred in seven samples, four of which were breast cancers (**Figure 3D**), and another rearrangement involving *CTTN* (this time connected to *ACTN3*) was detected in five samples, including three breast cancers. Both rearrangements were associated with increases in *CTTN* expression (1.9-fold and 4.9-fold, respectively) relative to lineage controls (**Supplementary Figure 3**B; combined p=0.03). CTTN (cortactin) interacts with cytoskeletal and cell adhesion complexes and has been hypothesized to drive 11q13 amplifications, which occur in 15% of breast cancers [35]. *CTTN* is within 1 MB of *CCND1*, a frequently amplified oncogene, and *CCND1* is also overexpressed in samples with the *CTTN* rearrangements (by 3-fold and 2.75-fold, respectively); thus these clusters may reflect consistent rearrangement patterns that give rise to *CCND1* amplicons. We conclude that further study is warranted into the effects of *GATA6* on GI cancers and into the roles of *CTTN* rearrangements on breast cancers, either by direct *CTTN* overexpression or shaping the contents of the *CCND1* amplicon.

### Accounting for Complex Connections

#### Connections Resulting from Combined Rearrangements

Across the entire sample set, 86% of SV events are simple (comprising a single rearrangement); the remaining 14% of events are complex, comprising multiple rearrangements--up to hundreds or more in the case of chromothripsis, breakage-fusion-bridge, or chromoplexy events [10,36,37]. As a result, 51% of rearrangements result from complex events (**Figure 4b**). These complex events can generate similarly complex topologies, in which distant genomic loci may be juxtaposed through the combined effects of two or more rearrangements (**Figure 4a**). For example, a “second-order connection” connects loci through two neighboring rearrangements; a “third-order connection” connects them through three.

**Figure 4.**
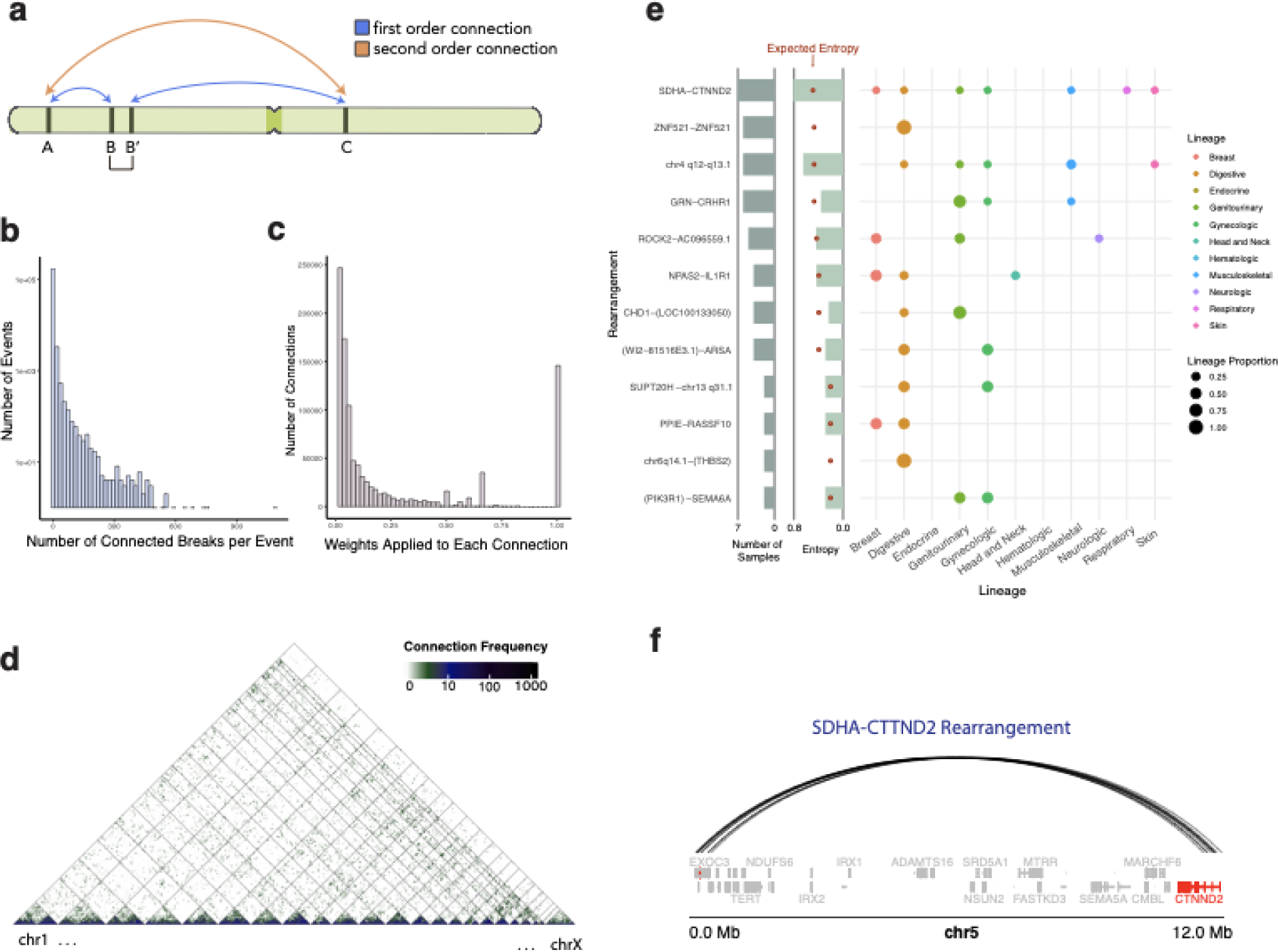
Significantly recurrent connections through more than one rearrangement as identified by SVSig-2Dc. a) Schematic of a “second-order” connection between A and C, through “first-order” connections between A and B and B’ and C. b) Distribution of the number of individual connections within each complex rearrangement event. c) Distribution of weights given to each connection across all rearrangements, including complex events. d) Observed connection frequencies genome-wide, including second- and higher-order connections, after applying weights in (c). Connection frequencies are represented on a log10 scale. e) Novel significant connections identified by SVSig-2Dc. For each event, the proportion of affected samples that fall under each lineage is shown by the size of the circle. Entropy values represent levels of lineage specificity. The number of samples containing each rearrangement is also indicated. f) A significant complex event connecting SDHA and CTTND2 in seven samples. The simple rearrangements that give rise to these connections are shown in Supplementary Figure X.

We were interested to understand how these higher-order connections, and complex events more generally, contribute to recurrent connections in cancer. To approach this, we developed SVsig-2Dc, a method that first detects all simple and higher-order connections, and then includes these in both the matrix of observed connections and the background model. To this end, we applied JaBbA [38] to determine the somatic distances between all breakpoints, and then classified all breakpoints within 1 Mb of somatic distance (approximately the size of a large TAD) as “connected” (**Figure 4a-b**). This required that we downweight individual connections within complex events, so as not to double-count the rearrangements they comprise.. In brief, each connection was down-weighted by the total number of connections within the connected component it belonged to (**Figure 4c**, see Methods). For example, a complex event comprising two adjacent rearrangements would have three connections; thus, each connection counts as ⅔ of a rearrangement (**Figure 4a**). Using this method, we detected 1,031,832 connections overall, of which 86% involved two or more rearrangements. On average, connections within complex events involved approximately three (3.14; sd 10.75) genomic loci (**Figure 4b**).

### New Candidate Driver Rearrangements

Surprisingly, accounting for higher-order connections reduced the number of significantly recurrent connections from 80 in the SVsig-2D analysis to 29 in SVsig-2Dc (Supp. Table 7). This is because first-order connections within complex events were down-weighted to account for connections involving two or more rearrangements. The 29 significant connections included 17 that had been detected by SVsig-2D and 12 novel events, of which 6 reflected at least some higher-order connections. Although the total number of unique events in SVSig-2Dc was much lower, more samples were implicated in each recurrent event (an average of 11 per event, vs 5 per event in SVSig-2D).

SVSig-2Dc identified the same top hits as SVSig-2D and introduced new candidate driver events. All events that occurred across 10 or more samples were detected by both models. The new events identified by SVSig-2Dc involved two to seven samples per hit. Furthermore, all new events were intrachromosomal, and overall, SVSig-2Dc showed a slight preference for intrachromosomal events (17/29 detected) compared to SVSig-2D (41/80 detected). Again, these events were highly tissue type specific; 90% of hits comprised a majority of connections from one lineage vs 26% in random data (permutation test, p<1e-4). Most complex events (17/29) also experienced lower entropy within each cluster than expected by chance, though this association was weaker than in the SVSig-2D analysis (**Figure 4e**).

Among the novel events, a *SDHA-CTNND2* rearrangement was most significant. This was present across seven samples from distinct subtypes and comprised complex events in two of those samples (**Figure 4f**). SDHA is a mitochondrial enzyme involved in energy conversion, and germline alterations in SDHA have been previously implicated to predispose individuals to multiple cancers, including paraganglioma, pheochromocytoma, neuroblastoma, and gastrointestinal stromal tumors [39,40].

## Discussion

The distribution of rearrangements across the cancer genome reflects biases introduced by the mechanisms that formed them and the effects of evolutionary selection, promoting events that increase cellular fitness and decreasing observed frequencies of events that harm cellular fitness. The frequency of rearrangements is strongly shaped by sequence, epigenetic, and spatial features, and is most predicted by the span of the rearrangement. All of these likely reflect mechanistic biases. Junction microhomology is known to correspond to the DNA damage repair mechanisms that formed rearrangements [23,31,41], and we found that the implied repair mechanisms were differentially enriched across known cancer genes and tumor subtypes.

SVSig-2D makes use of these predictive features of rearrangements to create a background model of expected rates at which pairs of loci will be connected by rearrangements in the absence of selection. It then detects candidate positively selected “driver” rearrangements as those that recur surprisingly often given this background rate. Similar methods have been previously developed to detect significant SNVs, CNVs, and breakpoints, ([7], [17], [8]). However, all of these look for recurrent events within the one-dimensional continuum of the genome. SVSig-2D is the first significance method to detect candidate driver events in a two-dimensional space–in this case, connections between pairs of genomic loci–after controlling for the frequency of breaks at each of the involved loci. SVSig-2Dc, which accounts for complex connections, effectively makes use of higher-order dimensions of rearrangements. This affects the statistical power that can be reached, because larger cohort sizes are required to adequately sample higher-dimensional spaces(ref). Recent decreases in the cost of sequencing, and associated increases in the number of tumors undergoing whole-genome sequencing(refs) will be useful to improve our ability to detect rare driver rearrangements.

Like all methods that detect candidate driver events based upon their recurrence rates, SVSig-2D and SVSig-2Dc can provide false positive or negative results if their background models are incorrect. Thus, special care must be taken in generating background models, and all candidate driver events detected by these methods require experimental validation.

Even so, SVSig-2D and SVSig-2Dc introduced a new framework for differentiating background and driver rearrangements, and further possibilities exist for improving the method. Future iterations of this method may incorporate additional rearrangement features, such as the topologies in which genomic loci join, or sequence features such as homologies between breakpoints. Additionally, we may try to better characterize complex events. One approach may involve extracting relevant information from complex rearrangement signatures. Another approach may account for specific classes of rearrangements ([38]), or events such as chromothripsis and chromoplexy. Finally, information derived from the three-dimensional spatial organization of the genome may underscore rearrangements that are located in active chromatin regions, disrupt TADs, or alter likelihood of contact between poorly expressed genes and active enhancers.

## Methods

### Data Sources

The rearrangements analyzed in this paper were called from 2,693 WGS cancers sequenced by the ICGC and TCGA PCAWG Consortium. In brief, rearrangements were called from four separate callers and merged to generate high confidence consensus calls. Details on the consensus calling are located in [16]. The final data can be found at https://dcc.icgc.org/releases/PCAWG/consensus_sv. Cancer types were grouped into lineage classifications based on the ICD10 [42] and NCI cancer type groupings (https://www.cancer.gov/types/by-body-location). Gene expression data were obtained from https://dcc.icgc.org/releases/PCAWG/transcriptome/gene_expression, and sequence features (epigenetic tracks, fragile sites, TADs, and LTR, LINE, and SINE regions) were collected from data sources as described in Li et al, 2020.

### Calculating a 2D background model of rearrangement frequency

We represented the distribution of rearrangements within an adjacency matrix, where each dimension of the matrix, which we denote as *i* and *j,* represents the 1-dimensional genome split into approximately 500kb bins. The values in the matrix represent the number of rearrangements that have breakpoints within both bins *i* and *j*.

We then calculated two background models to represent these data. Each background model (double-break join and break-invasion) determines the probability of a rearrangement occurring in the *ij*th tile and is solved separately. The double-break join probability is described in Equation 2.

*r_i_* and *r_j_* are the probabilities of a break in locus *i* and *j*, respectively, and *l_ij_*is a length factor that represents the distribution of observed rearrangement spans.

In contrast, the break-invasion probability is represented in Equation 1.

where *q_i_*is related to the probability of a break in locus *i* and *s_ij_*is the conditional probability of an invasion into locus *j* given a break in locus *i*. Since it is not possible to distinguish the broken locus from the invaded locus, the overall probability *p_ij_* is the sum of both reciprocal terms. The terms *q* and *r*, and *s* and *l*, are normalized to keep units constant. In the break-invasion model, *s_ij_*is *l_ij_*, normalized such that the sum of probabilities *r_i_ s_ij_* over all *i* equals one, and *q_i_* is determined by solving the equation *(s_ij_ + I) q_i_ = r_i_*, where *I* represents the identity matrix.

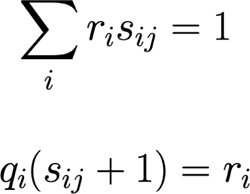

The final background model is a linear combination of the break-invasion and double break join models (Equation 3).

The weight between these two models (denoted as α) is solved by finding the best fit of the background model to the data by minimizing the Bayesian Information Criterion (BIC). This alpha parameter was calculated separately for short (500kb-1Mb), long (larger than 1Mb), and interchromosomal tiles.

### Accounting for complex juxtapositions

To account for rearrangements that occur in proximity to each other, we grouped nearby rearrangements and downweighted their individual contributions to sum to a single complex rearrangement. Specifically, for every pair of rearrangements with breakpoints within 1 Mb of somatic distance, we defined both the region in between those rearrangements and the distant breakpoints to which they connect as “connected components”. To avoid overcounting complex rearrangements with many such connections, we weighted each first- and higher-order connection according to the number of simple “first-order” connections involving the connected component, divided by the total number of connections involving that connected component. This could result in substantial down-weighting of individual rearrangements. For example, a complex event involving 10 rearrangements could result in 55 (11 choose 2) first- and higher-order connections, and each of these connections could be downweighted by a factor of 5.5. In effect, this approach allows for an increased number of connections but decreases the certainty associated with any one of them. Because the weights used by SVSig-2Dc were not always integers, we used a continuous generalization of the binomial distribution[43] to calculate probability distributions for the number of rearrangements per tile.

### Identifying significantly recurrent rearrangements

Significance for each tile was determined according to both the enrichment rearrangements within the tile and how tightly these events clustered. We first used the cumulative binomial distribution (*N*, *p_ij_*) to calculate the probability (p-value) of observing N or more events in that tile. Next, we calculated the 2D distances between every combination of events within the tile, and determined the probability (p-value) that the mean of this distribution, or a smaller mean, would be observed by chance. To do so we compared this observed distribution to the expected distribution from simulated random events within that tile using a Student’s t-test. We combined these rearrangement frequency and clustering p-values with Fisher’s method and applied an FDR multiple hypothesis correction to determine the final significance (q-value) for each tile.

Once we identified putative driver rearrangements, we removed any candidate rearrangements that occurred in only one sample or contained potential artifacts. Primarily, we removed rearrangements where breakpoint clusters had breakpoints within a standard deviation of 10 base pairs, out of concern that these rearrangements were likely to be sequencing artifacts or mis-called germline events. The remaining rearrangements were annotated according to genes proximal to its breakpoints. Rearrangements were annotated according to the most proximal genes to both breakpoints, with two exceptions. When a known COSMIC cancer gene was within 500 kb of a breakpoint, the COSMIC gene was used for annotation, and when multiple genes occurred within 10 kb of a breakpoint, the alphabetically first gene was used.

### Simulating rearrangement distributions

To test the capacity of the model to distinguish between known rearrangement distributions, we simulated rearrangements following pre-determined background models. We took linear combinations of the break-invasion and double-break join models with α parameters of 0, 0.25, 0.5, 0.75, and 1. For each tile in the background model, we simulated the total number of events based on the binomial distribution (*N*, *p_ij_*), where *N* represented the total number of rearrangements across all tiles and *p_ij_* was the expected probability of a rearrangement occurring within that tile per the background distribution. Within each tile, we then randomly assigned a “start” and “end” breakpoint to each simulated event.

To test the likelihood with which the model detects false positives, we simulated rearrangements without positive selection. We preserved the span distribution and marginal breakpoint distributions for intrachromosomal tiles. For every 1Mb bin of the span distribution, we calculated the percentage of events that would fall in each tile, based on the area of the tile accounted for by the span bin. Thus, sections of each tile corresponded to a different range of spans. We then multiplied each tile section by the sum of its marginal breakpoint distributions, ri and rj, to obtain an expected probability of events, *p_ij,_*. We then calculated the number of events within each tile section based on the binomial distribution (N*_intra,_ p_ij_*), where *N* represented the total number of intrachromosomal rearrangements. Finally, we sampled spans for each event from its corresponding span distribution and assigned start and end coordinates within the tile. For interchromosomal tiles where a span could not be calculated, the number of events per tile were simulated based on the binomial distribution (N*_inter,_ p_ij_*), where N*_inter_* represented the total number of interchromosomal rearrangements. *p_ij_* represented the expected probability of interchromosomal rearrangements within that tile, with correction for the marginals.

### DNA Damage Repair Clustering Analyses

Rearrangements were assigned to a DSB repair mechanism based on the length of microhomology present at their breakpoints. Rearrangements with 0, 4 - 9 bases, or 13 or more bases of microhomology were assigned to NHEJ, MMEJ, and SSA, respectively. Rearrangements with 1-3 and 10-12 bases of homology were designated “no call”. We then performed k-means clustering to group samples based on the total number of rearrangements within that sample and the proportion of called rearrangements assigned to each DSB repair mechanism. We assessed enrichment for tissue types and known driver genes (https://dcc.icgc.org/api/v1/download?fn=/PCAWG/driver_mutations/TableS3_panorama_driver_mutations_ICGC_samples.public.tsv.gz) within each cluster by performing a Fisher’s exact test, denoting significant tissues when p<0.05. An enrichment effect size was derived from the Fisher’s test statistic. Specific alterations to each gene were annotated, including coding/noncoding status, and if the gene was amplified, deleted, or fused.

### Lineage enrichment of significant events from SVSig-2D and SVSig-2Dc

The propensity for significant hits to fall under a single lineage was determined with entropy calculations. In brief, individual samples were assigned to a lineage as described above. Next, for each event cluster, we calculated the number of samples in that cluster belonging to each lineage. We calculated the entropy, H(x) for each significant hit *x* following Eq. 6, where *n* represents the total number of unique lineages found within that hit, and b is the total number of lineages found across all hits.

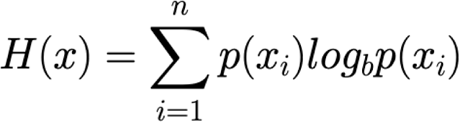

To assess the level of non-randomness present in the lineage distributions across significant hits, we performed permutation tests. We assigned samples randomly to a cluster while preserving the total number of samples from each lineage and within each cluster. We then calculated the number of clusters that had more than half of its samples derived from a single lineage. We compared this distribution to the true number of clusters that were made up of a majority of one lineage.

## Code Availability

The code to detect significantly recurrent simple and complex rearrangements with SVSig-2D and SVSig-2Dc are freely available for use at https://github.com/beroukhim-lab/SVsig.

---

**Supplementary Figure 1.**
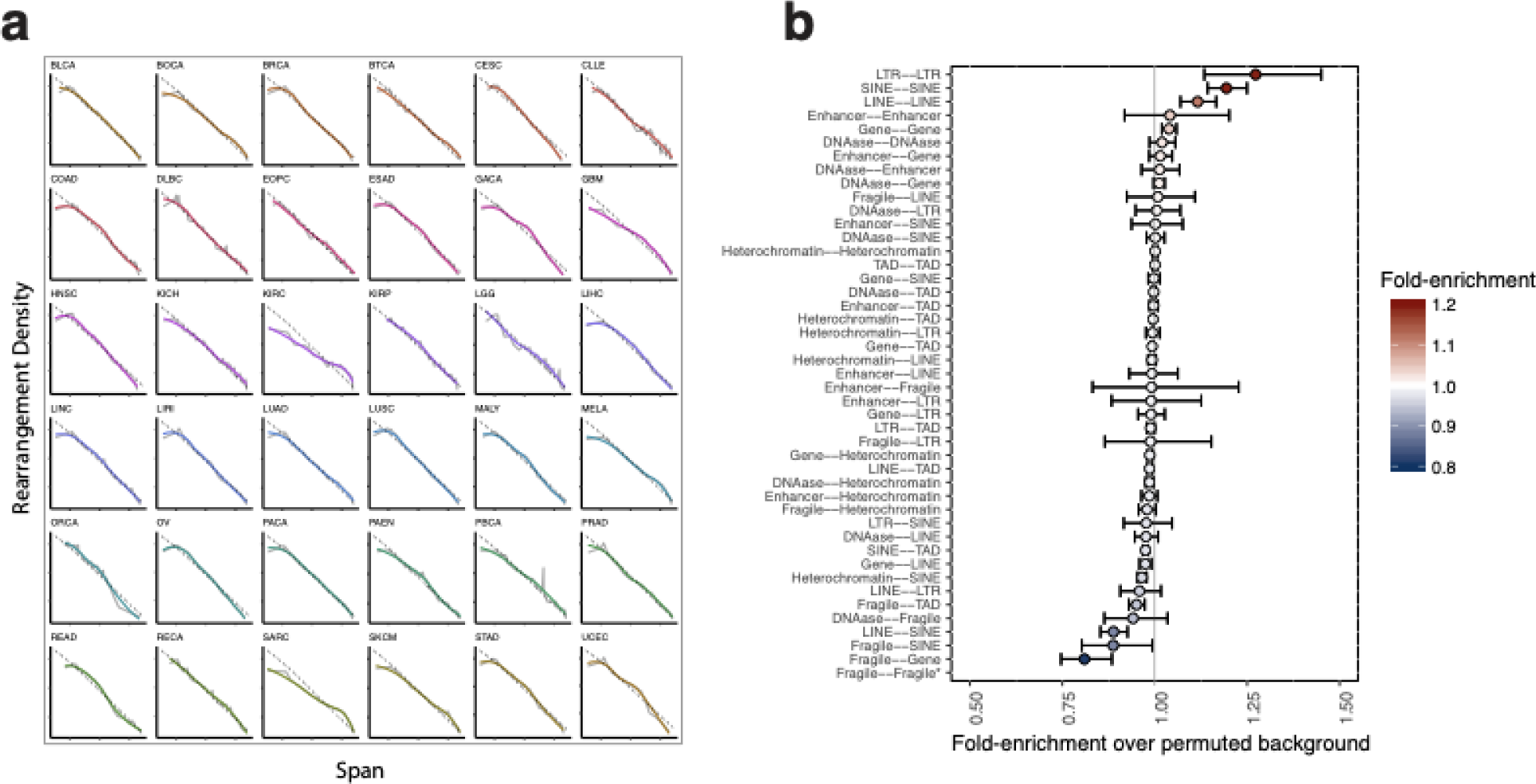
Additional features predictive of rearrangement formation. a) Rearrangement spans, separated by tumor type. The 1/span distribution is represented in gray. Tumor types are labeled by their corresponding ICGC project code. b) Connection frequencies across epigenetic and sequence features.

**Supplementary Figure 2.**
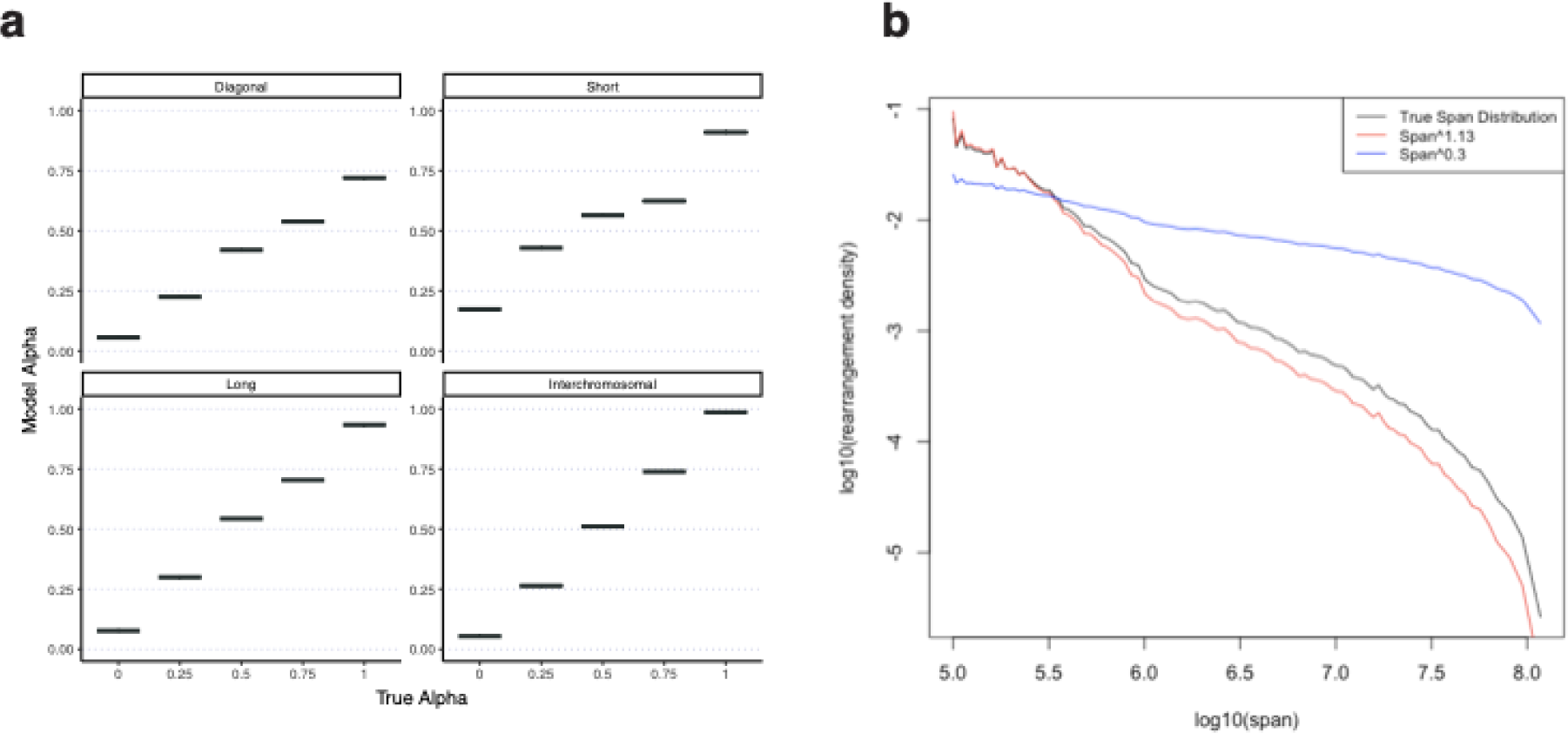
Stability of the background model. **a)** Model predictions of α values for diagonal, short (<1Mb), long intrachromsomal (>1Mb), and interchromsomal tiles (vertical axis), using simulated data with actual α values shown on the horizontal axis. **b)** New span distributions after raising the true span distribution by a range of powers. The model produced false positives when the span distribution was raised to a power less than 0.3 (blue) or greater than 1.13 (red).

**Supplementary Figure 3.**
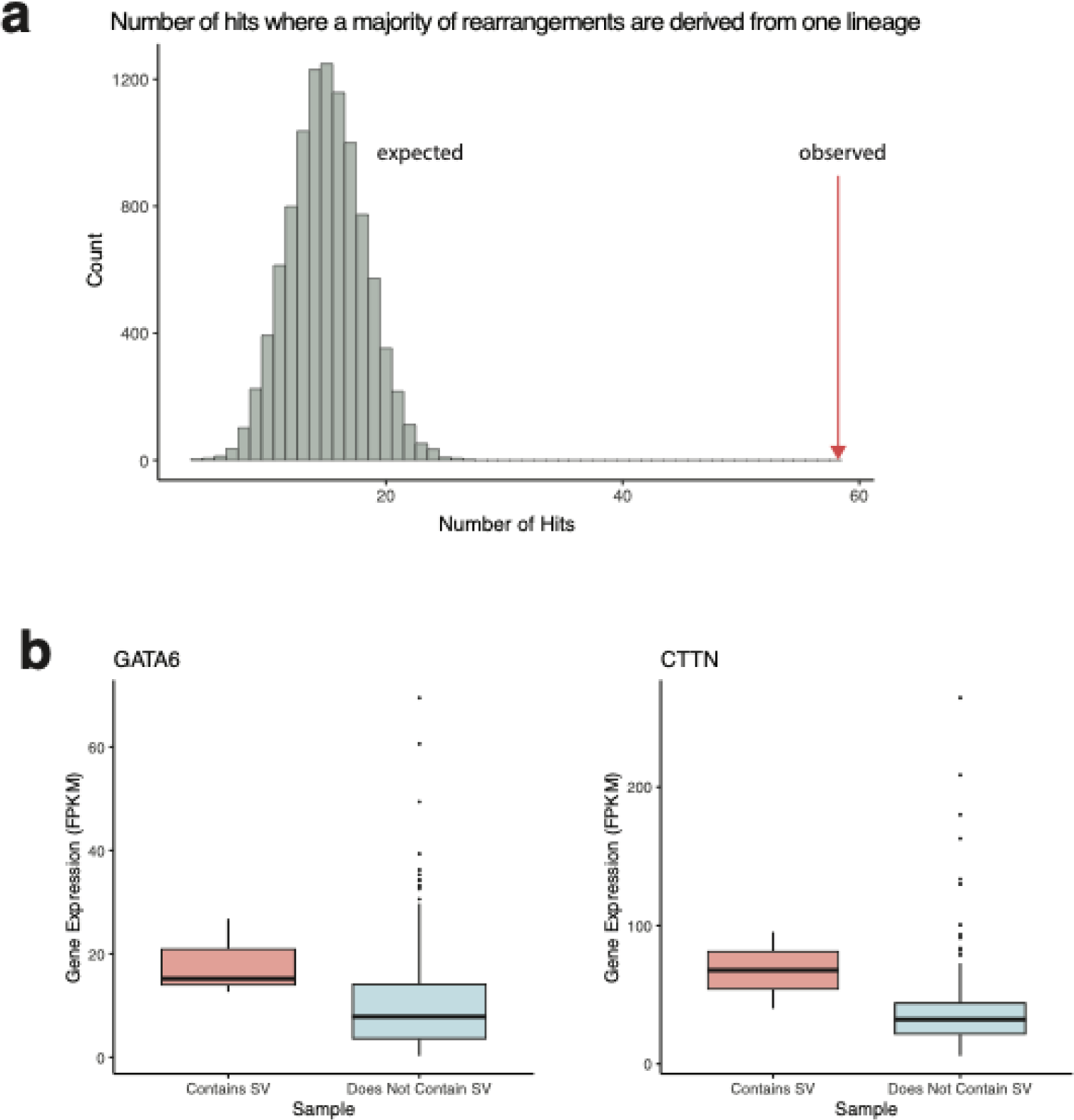
Significant recurrent rearrangements from SVSig-2D are lineage-specific and associated with changes in gene expression. a) Observed and expected number of significant events where the majority of samples with the event derived from the same lineage. The expected distribution was generated by permuting samples across all significant events, preserving the number of lineage types and number of samples per event. b) Gene expression (FPKM) from RNA-seq of GATA6 for samples with and without significantly recurrent rearrangements in GATA6 (p=0.06). c) Gene expression (FPKM) from RNA-seq of CTTN for samples with and without significantly recurrent rearrangements in CTTN (p=0.03).

